# *Fusarium graminearum-*induced shoot elongation and root reduction in maize seedlings correlate with later seedling blight severity

**DOI:** 10.1101/287714

**Authors:** Shaoqun Zhou, Justin S. Bae, Gary C. Bergstrom, Georg Jander

**Affiliations:** Boyce Thompson Institute, 533 Tower Road, Ithaca, NY 14853; Plant Biology Section, School of Integrated Plant Science, Cornell University, Ithaca, NY 14853; Plant Pathology and Plant-Microbe Biology Section, School of Integrated Plant Science, Cornell University, Ithaca, NY 14853

**Keywords:** *Fusarium graminearum* seedling blight, maize, shoot elongation, rootreduction.

## Abstract

*Fusarium graminearum* seedling blight is a common disease of maize (*Zea mays*). Development of genetic resistance to seedling blight in maize germplasm requires efficient and accurate quantitative assessment of disease severity. Through artificial inoculation experiments under controlled growth conditions, we determined that host genotype, pathogen genotype, and infection dose influence the extent to which *F. graminearum* induces shoot elongation and inhibits root growth in maize seedlings. A comparison of fifteen maize inbred lines showed independent variation of these two fungus-induced effects on seedling growth. In a broader survey with nine commercial maize hybrids and three field-collected fungal isolates, there was significant correlation between these seedling growth responses, as well as with later seedling blight severity. Analysis of variance suggested that this variation and the observed correlative relationships were primarily driven by differing pathogenicity of the three fungal isolates. Together, our results indicate that *F. graminearum*-induced shoot elongation and root reduction in maize seedlings have distinct underlying physiological mechanisms, and that early observations of seedling growth responses can serve as a proxy for investigating natural variation in host resistance and pathogen aggressiveness at later growth stages.

## INTRODUCTION

*Fusarium graminearum* Schwabe (teleomorph: *Gibberella zeae*) is a widespread fungal phytopathogen that causes seedling blight, root rot, stem rot, or ear rot in maize (*Zea mays*), depending on the timing and tissue of infection (Munkvold and White 2016). The severity of maize diseases caused by *F. graminearum* in the United States varies significantly from year to year, causing 15 to 80 million bushels of annual yield loss due to ear rot, and another 30 to 85 million bushels due to stalk rot (Mueller 2016a, b, c, 2017). Unlike Gibberella ear rot and stem rot diseases caused by *F. graminearum*, which can be tracked by the characteristic pink-reddish pigmentation of fungal mycelium, seedling blight and root rot diseases are often caused by a consortium of soil-dwelling fungal and oomycete pathogens, including *F. graminearum*. While the dominant species/strains of these consortia are likely to vary considerably across geographic locations, collectively these seedling blight and root rot diseases can account for over 200 million bushels of annual yield loss, making them among the most damaging maize diseases in the northern states (Mueller 2016b). Management of *F. graminearum* seedling blight can be difficult because standard seed fungicide treatment is not always sufficient and may promote the development of resistance in the fungal population. Genetic sources of seedling blight resistance have yet to be established in commercial maize germplasm.

In addition to causing direct yield loss through tissue rotting, *F. graminearum* can also cause crop contamination with mycotoxins such as deoxynivalenol and zearalenone, leaving the grain and stover unsafe for human and livestock consumption, and imposing further financial burdens on the growers (Arunachalam and Doohan 2013; Maresca 2013). Furthermore, *F. graminearum* can also cause destructive scab and head blight diseases in barley and wheat (McMullen et al. 2012; Trail 2009). Considering the overwinter persistence and airborne infectious ascospores of this pathogen, unchecked *F. graminearum* diseases in maize could have a far-reaching negative impact on other crop species in the vicinity.

Breeders and researchers interested in improving resistance against *F. graminearum* in maize have been doing artificial inoculation experiments in both field and controlled environments, primarily focused on infected maize ears and stalks. Genetic mapping of Gibberella ear rot resistance based on symptom severity has identified a large number of quantitative trait loci (QTL), each with small effect size. Moreover, these QTL tend to be sensitive to experimental methods and environmental conditions, making them difficult to validate across different studies (Ali et al. 2005; Brauner et al. 2017; Kebede et al. 2016). In contrast, major genetic loci contributing to Gibberella stalk rot resistance have been identified with bi-parental mapping populations (Chen et al. 2017; Ma et al. 2017; Yang et al. 2010; Zhang et al. 2012). Ye et al. (2013) observed that maize near-isogenic lines resistant against Gibberella stalk rot also showed less severe seedling root browning and shrinkage after *F. graminearum* inoculation. This suggests that resistance against Gibberella stalk rot and *F. graminearum* seedling blight, both diseases in vegetative tissues, are likely controlled by similar genetic and physiological mechanisms, and are distinct from those affecting Gibberella ear rot resistance. This is consistent with the known influence of tissue type and developmental stage on the interactions between maize and *F. graminearum* (Zhang et al. 2016).

Genetic improvement of *F. graminearum* resistance depends on accurate quantitative assessment of resistant and susceptible phenotypes. Here we show that *F. graminearum* inoculation of maize seedlings not only inhibits root growth, but can also promote shoot elongation in a dosage- and genotype-dependent manner. Shoot elongation in response to *F. graminearum* root infection is significantly correlated with seedling survival, suggesting that these traits could be used to assess seedling resistance, as well as serving as a quantitative measure of the aggressiveness of different *F. graminearum* isolates.

## MATERIALS AND METHODS

### Plant and fungal materials

Fifteen parental lines from the maize nested association mapping population (McMullen et al. 2009) were chosen for *F. graminearum* infection assays. For a broader survey, nine organic commercial maize hybrids were kindly provided by the Blue River Organic Seed Company (Table 1). The maize hybrids were selected to cover different maturation time and provider-assessed seedling vigor levels. The accession numbers of the tested hybrids are masked per the provider’s request. Three field-collected *F. graminearum* isolates (Table 2) that have shown different levels of aggressiveness in a previous laboratory test (Kuhnem et al. 2015) were used for inoculations (Table 2). Fungal chemotype, namely the type of major mycotoxin accumulated, was not considered during isolate selection, since it was not significantly associated with aggressiveness in the previous test.

### Plant growth and fungal inoculation methods

Maize seeds were germinated in moisturized rolls of germination paper after surface sterilization with 10% household bleach for 30 minutes. Germinated seedlings were transplanted to 7.5 cm × 7.5 cm plastic pots with Turface® MVP® calcined clay (Profile Products LLC, Buffalo Grove, IL) when their primary roots reached approximately 9 cm, which would take 3-5 days depending on the specific maize genotype. For inoculation experiments, seedling roots were immersed in *F. graminearum* spore suspension or mock solution (0.03% Phytagar) for one hour prior to transplanting. Fungal spore suspensions were freshly prepared by flooding and scraping 7-day-old fungal cultures maintained on Potato Dextrose Agar plates under black lights with 0.03% Phytagar suspension. The spore concentration was estimated with a hemocytometer (Hausser Scientific, Horsham, PA) and light microscopy. Fungal hyphae fragments were observed in spore suspension by light microscopy.

For the concentration gradient experiment, twenty-four seedlings of each maize hybrid were grown, and six were inoculated with each tested fungal spore concentration and mock inoculum. For each pair of maize hybrid and fungal isolate included in the diversity screening, at least 15 seedlings were inoculated with fungal spores, and 5 were mock-inoculated. Transplanted seedlings were kept at 26°C (Day)/23°C (Night), 60% humidity, and a 16 hour long-day light cycle. Seedlings inoculated with different *F. graminearum* isolates or mock inoculum were kept in separated growth chambers set to identical growth conditions.

### Phenotypic data collection and analysis

The heights of fungus- and mock-inoculated seedlings were measured nine days post-inoculation. On the same day, seedlings were removed from pots to measure their total root length using an image analysis algorithm implemented in the RootReader 2D software (Clark et al. 2013). After measurement, seedlings were re-potted in the same Turface particles. At fourteen days post-inoculation, the number of surviving seedlings within each fungus-inoculated maize hybrid population was counted as a measurement of *F. graminearum* seedling blight severity. Seedlings were considered dead when they broke at the soil line, with browning and rot at the breakpoint.

For seedling height and total root length, an average ratio of mock- and fungus-inoculated seedlings for each maize hybrid-fungus isolate combination was calculated to shown variation in these fungus-induced responses. Measurements from fungus- and mock-inoculated seedlings were compared using Student’s *t*-tests. Average ratios were rounded to one when measurements from the two groups were not significantly different from one another. For two-way ANOVA, maturation times of maize hybrids were grouped into four bins, as indicated in parentheses in Table 1. The average ratios of seedling height and total root length between fungus- and mock-treated seedlings were used as quantitative measurements to calculate linear correlation with seedling survival rate measured at 14 days after infection.

## RESULTS

### *Fusarium graminearum* inoculation at low concentration induces shoot elongation and root reduction in maize seedlings

Different concentrations of spore suspensions have been used to study natural variation in maize seedling responses to *F. graminearum* inoculation (Ye et al., 2013; Kuhnem et al., 2015). To determine the optimal concentration of *F. graminearum* spores that leads to the largest change in plant morphology, two concentration gradient experiments were performed with Gz014, a field-collected *F. graminearum* strain (Table 2), and two commercial maize hybrids with differing seedling vigor levels (Hybrids IV and V in Table 1, as assessed by the provider, Blue River Organic Seed Company).

At 1.2 × 10^6^ spores/mL, a concentration that is comparable to a previous study, the seedling stunting phenotype that had been reported was not observed in either of the tested maize hybrids (Kuhnem et al. 2015). Instead, the hybrid with high seedling vigor showed no significant change in seedling height, whereas the moderately vigorous hybrid seedlings grew significantly taller when inoculated with *F. graminearum* spores. At lower concentrations, fungus-induced shoot elongation became obvious in both highly and moderately vigorous hybrid seedlings (Figure 1A). Underground, *F. graminearum* infection significantly reduced root growth in the moderately vigorous hybrid, with no evidence of a dosage response. This fungus-induced change was not observed in the highly vigorous hybrid, which showed no significant change in total root length at all tested doses (Figure 1B).

**Figure 1.**
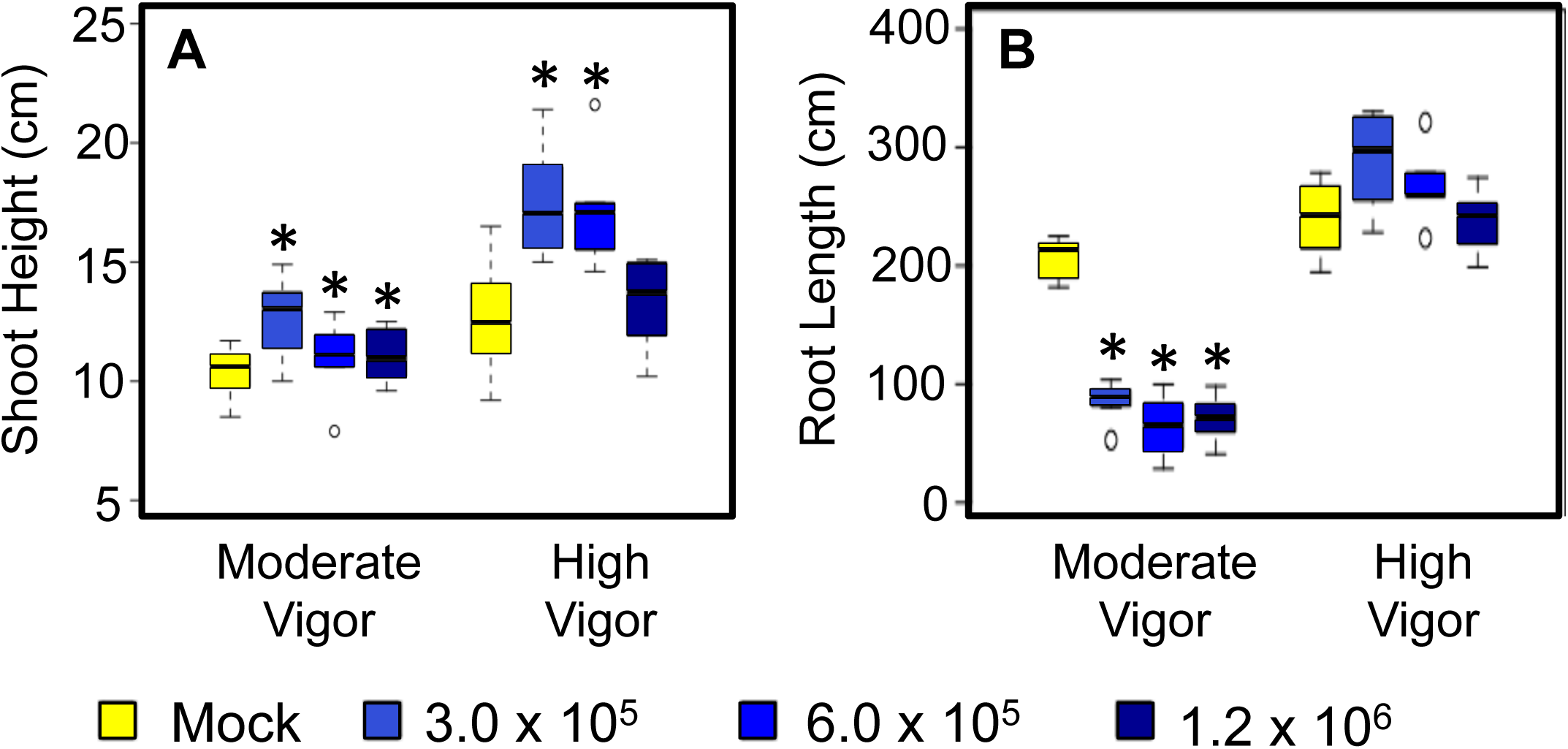
*Fusarium graminearum* inoculation induces shoot elongation and root reduction in maize seedlings in a dosage- and host genotype-dependent manner. Shoot height (A) and total root length (B) measurements on seedlings inoculated at each fungal spore concentration were compared to mock-treated seedlings with Dunnett’s tests relative to mock-infected controls (N = 6 for each group; * p < 0.05). Group means are indicated by the black bars, and the interquartile range by the upper and lower edges of the boxes. Outliers extending more than 1.5x of the interquartile range are indicated by individual circles, and the whiskers show the range of each group excluding the outliers. Fungal strain Gz014 was used for this experiment. All measurements were taken 9 days post-inoculation. Fungal spore concentration is in counts/milliliter.

For both fungus-induced growth responses, inoculation with the lowest fungal spore concentration (3 × 10^5^ count/mL) resulted in a response that was stronger than or comparable to that observed with the two higher inoculum concentrations. Therefore, the 3 × 10^5^ spores/mL *F. graminearum* dose was used for all subsequent experiments.

### *Fusarium graminearum*-induced shoot elongation and root reduction vary independently across maize inbred lines

To confirm the *F. graminearum-*induced shoot elongation and root reduction observed in the concentration gradient experiment, the artificial inoculation experiment was repeated with the reference maize genetic inbred line B73 with the same fungal isolate following the same protocol. Consistently, the same fungus-induced growth responses were observed (Figure 2). To investigate how *F. graminearum-*induced seedling shoot elongation and root reduction may be related, seedlings of fifteen maize inbred lines were inoculated with Gz014 or mock inoculum and their shoot and root growth were quantified 9 days after inoculation. Although *F. graminearum* inoculation reduced total root length in most inbred lines by approximately 50%, no significant difference was observed between mock- and *F. graminearum-* inoculated NC350, NC358, and Mo17 seedlings, (Figure 3A).

**Figure 2.**
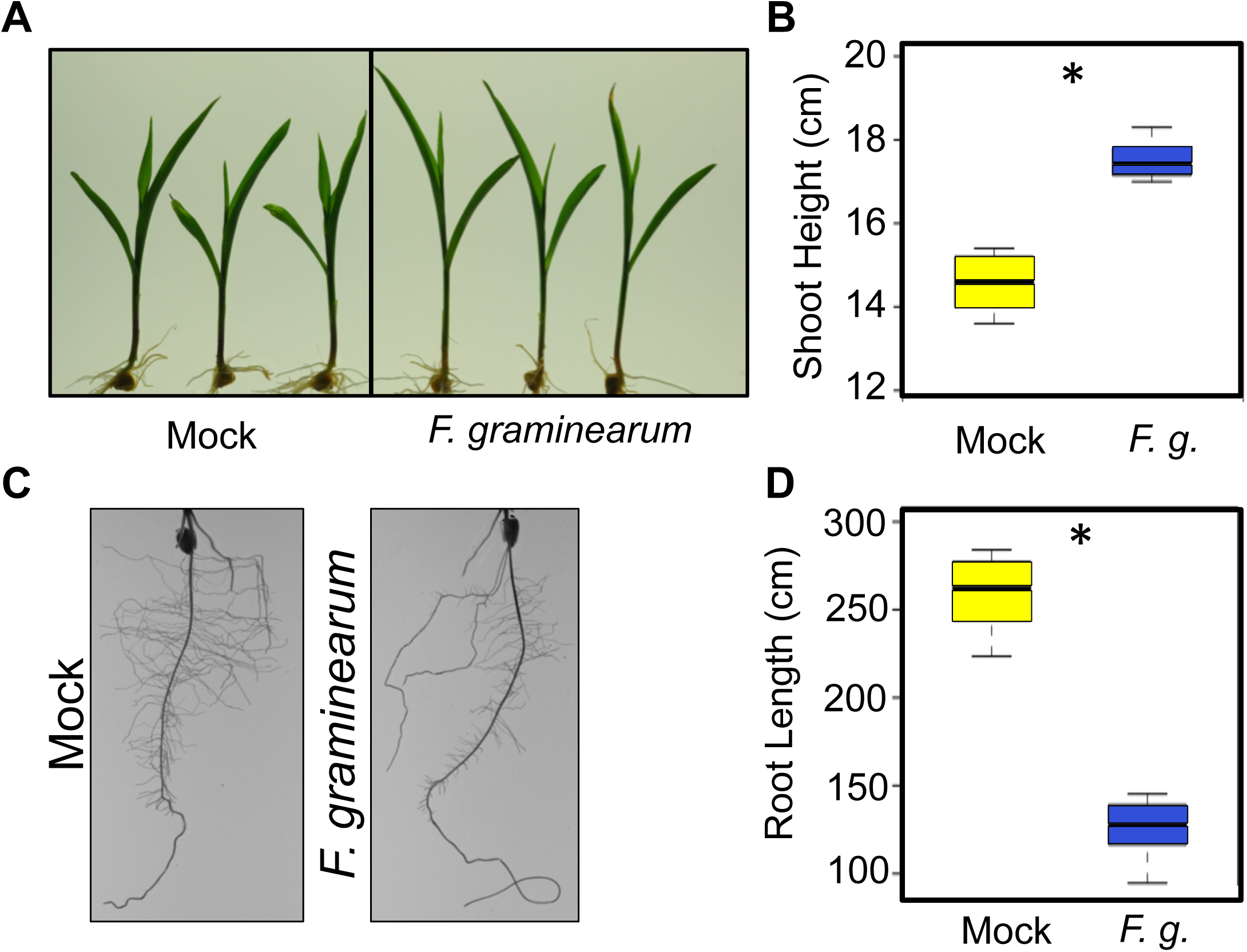
*Fusarium graminearum* inoculation induces shoot elongation and root reduction in maize inbred line B73. Representative photos of shoots (A) and roots (C) mock- and fungus-inoculated B73 seedlings were taken 9 days post-inoculation. Shoot height (B) and total root length (D) were measured on the same day and compared with Student’s *t-*tests (N = 6 for each group; *p < 0.05). Group means are indicated by the black bars, and the interquartile range by the upper and lower edges of the boxes. Outliers extending more than 1.5x of the interquartile range were indicated by individual circles, and the whiskers showed the range of each group excluding the outliers. Fungal strain Gz014 was used for this experiment. Spore concentration = 3 × 10^5^ counts/milliliter.

**Figure 3.**
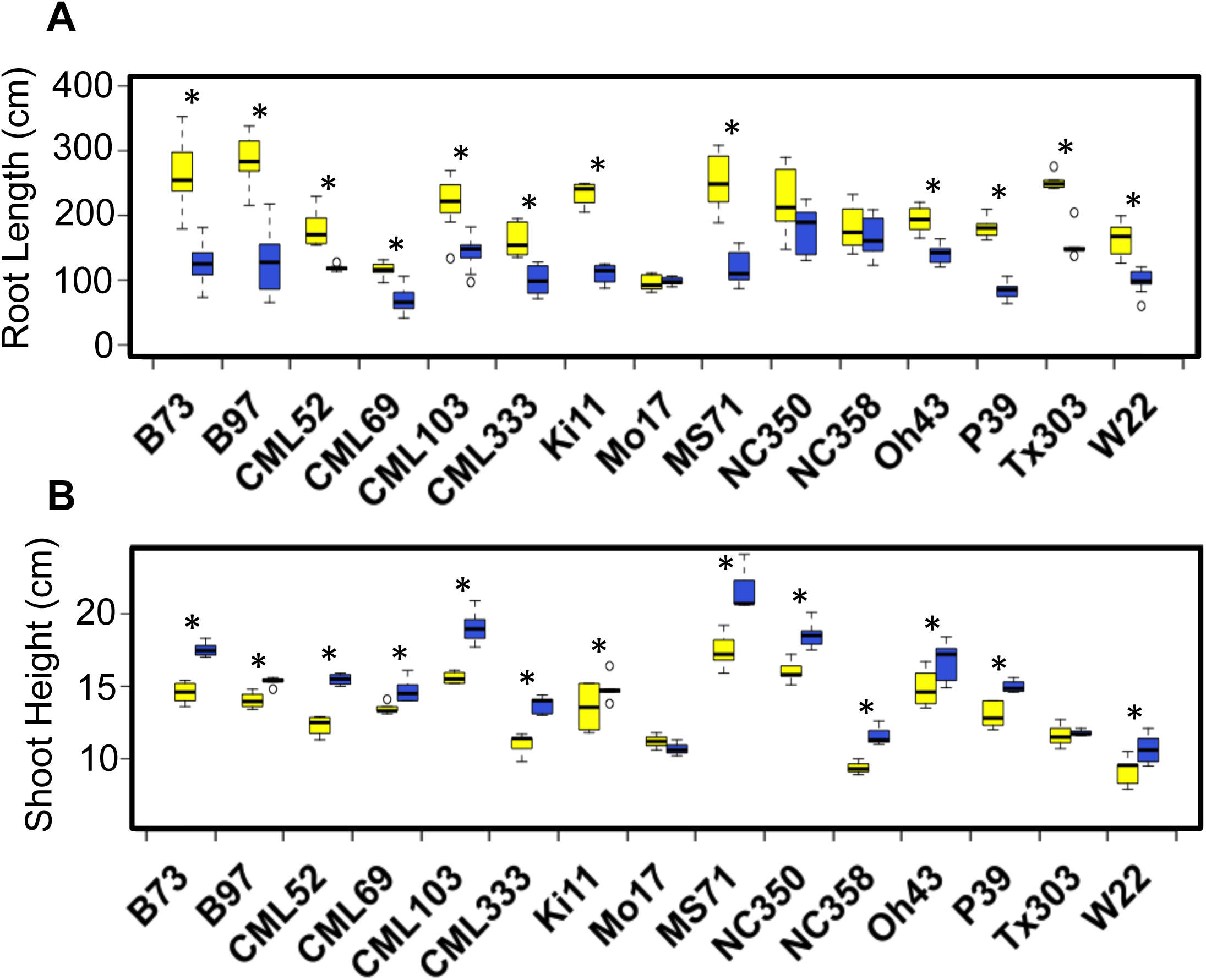
*Fusarium graminearum-*induced growth responses in maize seedlings vary across maize inbred lines. Root length (A) and shoot height (B) of each maize inbred line inoculated with *F. graminearum* spores (yellow) were compared to measurement from a corresponding mock treatment group (blue) with Student’s *t*-test (* p < 0.05). Group means are indicated by the black bars, and the interquartile range by the upper and lower edges of the boxes. Outliers extending more than 1.5x of the interquartile range are indicated by individual circles, and the whiskers show the range of each group excluding the outliers. All measurements were taken at 9 days post-inoculation. Fungal strain Gz014 was used for this experiment. Spore concentration = 3 × 10^5^ counts/milliliter.

In the same experiment, *F. graminearum*-induced shoot elongation also varied, ranging from 0 to over 25% increase (Figure 3B). Noticeably, the two fungus-induced growth responses do not correlate with one another across the fifteen tested inbred lines tested (Pearson’s R^2^ = 0.0079, p > 0.05). Two inbred lines showing no significant *F. graminearum*-induced root reduction, NC350 and NC358, grew 16% and 24% taller, respectively, after *F. graminearum* inoculation. Conversely, Tx303 only showed obvious root reduction (40%) but no significant shoot elongation (Figure 3). Inbred line Mo17 showed neither of the tested fungus-induced growth responses.

### *Fusarium graminearum-*induced growth responses and *F. graminearum* seedling blight severity are primarily determined by the fungal genotype

To further survey the natural variation in *F. graminearum*-induced seedling shoot elongation and root reduction, the artificial inoculation experiment was expanded to the nine commercial maize hybrids, including the two used in the concentration gradient experiment described above (Table 1), and three *F. graminearum* isolates (Table 2). Both *F. graminearum-*induced growth responses were highly variable across this panel (Figure 4). Two-way analysis of variance (ANOVA) performed on both traits showed that fungal isolate, rather than maize maturation time or provider-assessed seedling vigor, is the main source of assay variance (Table 3).

**Figure 4.**
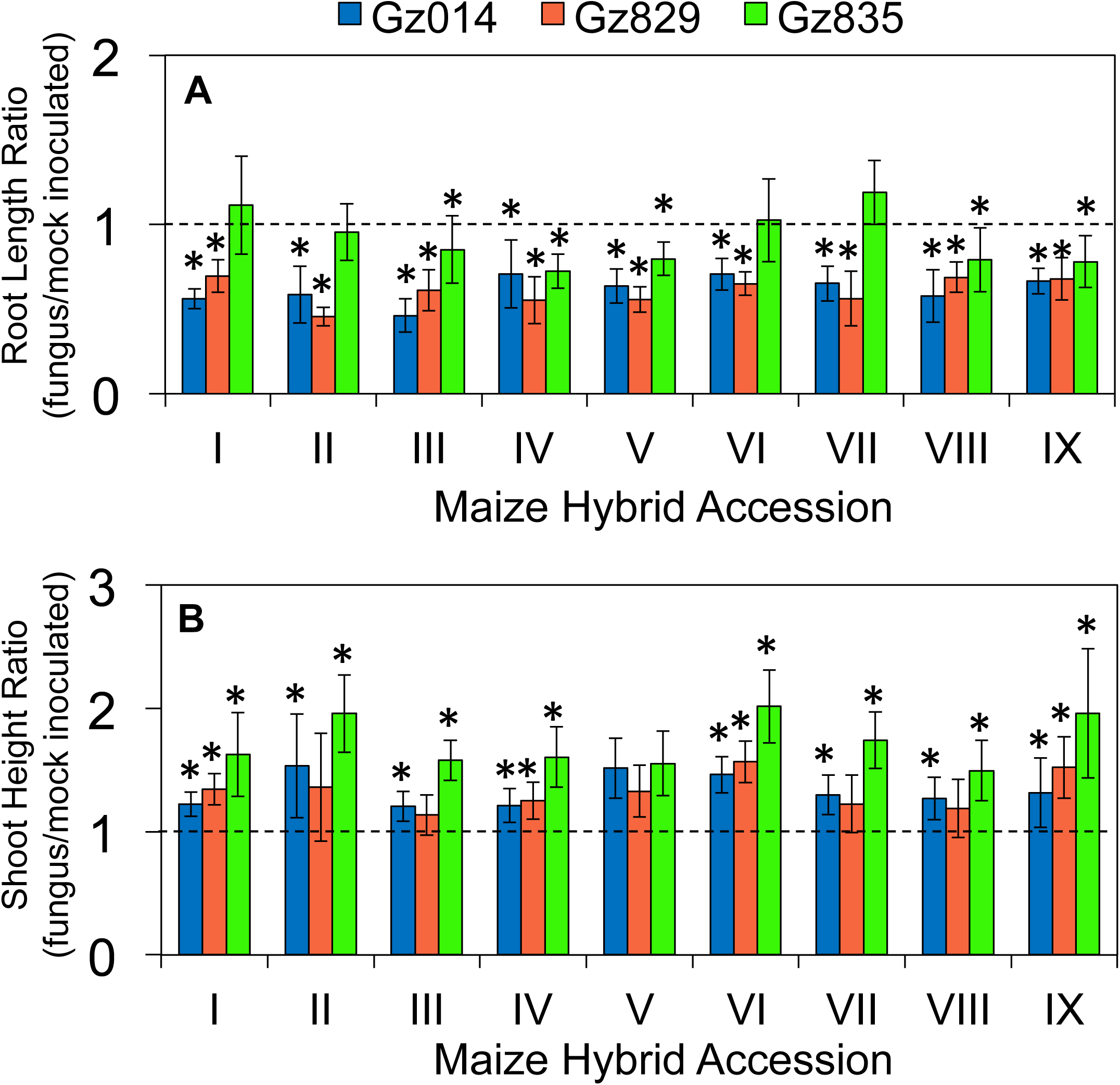
*Fusarium graminearum-*induced growth responses in maize seedlings vary with both plant and fungal genotype. Nine maize hybrid lines were inoculated with three *F. graminearum* isolates (Gz014, Gz829, and Gz835). Total root length (A) and shoot height (B) of each maize hybrid-fungus isolate pair were compared to measurement from a corresponding mock treatment group with Student’s *t-*test (* p < 0.05). All measurements were taken at 9 days post-inoculation. Fungal spore concentration = 3 × 10^5^ count/milliliter.

After quantifying *F. graminearum*-induced growth responses, the fungus-infected seedlings were kept under the same growth conditions, and *F. graminearum* seedling blight severity was quantified as the percentage of seedlings surviving in each maize hybrid population fourteen days after inoculation. As in the case of the morphological data, ANOVA of the seedling survival rate data identified the *F. graminearum* isolate as the most important contributor. Whereas both Gz014 and Gz829 infection left less than 50% of inoculated seedlings surviving after fourteen days, more than 85% of Gz835-inoculated seedlings survived in the same time period (Figure 5). The maturation time and seedling vigor of maize hybrids were not significant sources of variance (Table 3).

**Figure 5.**
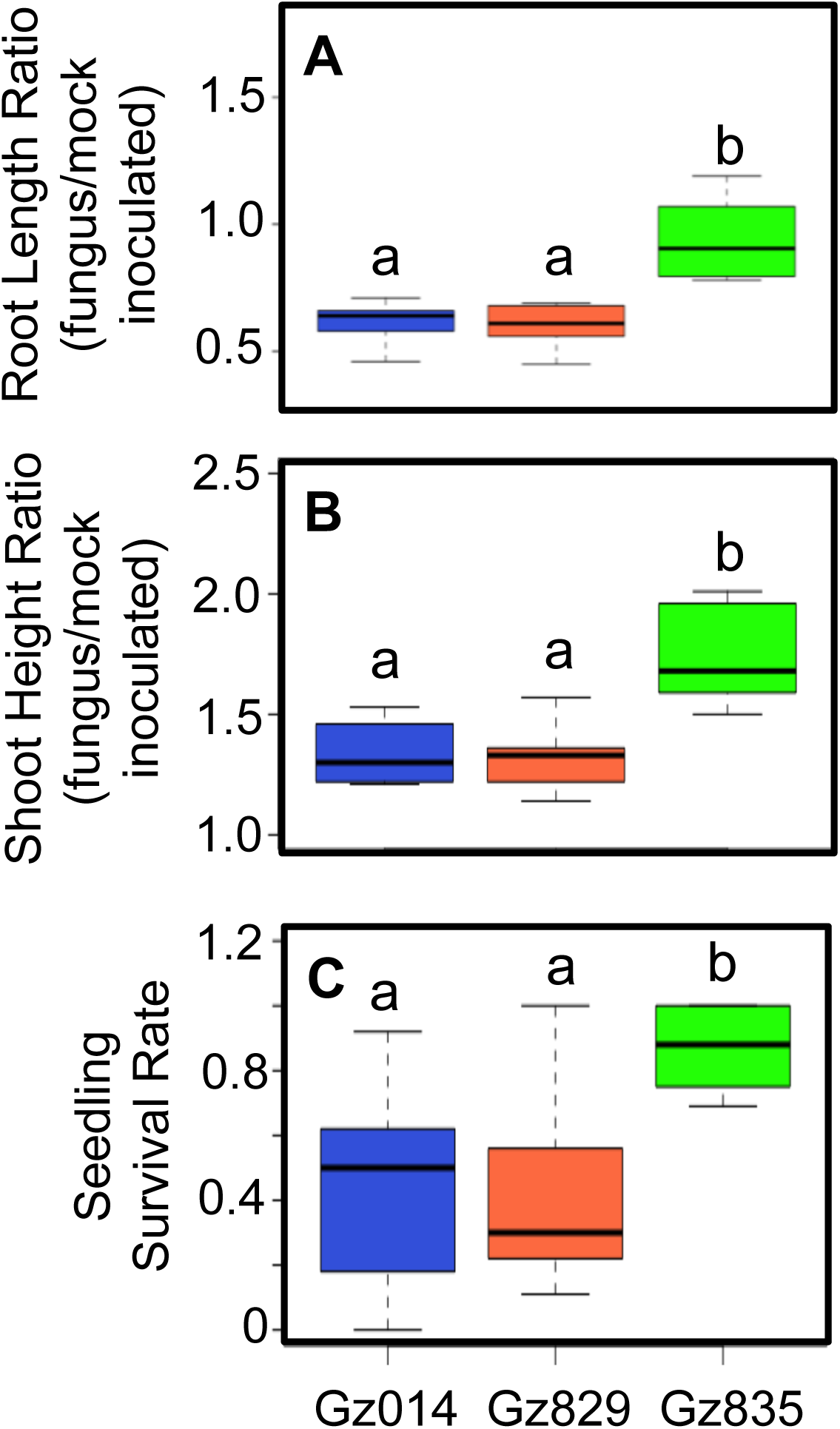
*Fusarium graminearum-*induced growth responses and seedling blight severity vary significantly between fungal isolates. The extent of *F.* graminearum-induced root reduction and shoot elongation at 9 days post-inoculation (A & B) and survival rates of maize seedlings inoculated with different *F. graminearum* isolates at 14 days post-inoculation (C) were compared using two-way analysis of variance with Tukey’s HSD. Group means are indicated by the black bars, and the interquartile range by the upper and lower edges of the boxes. Outliers extending more than 1.5x of the interquartile range were indicated by individual circles, and the whiskers showed the range of each group excluding the outliers. Significantly different groups are denoted with different letters. Fungal spore concentration = 3 × 10^5^ count/milliliter. Ratios refer to fungus/mock-inoculated plants.

### *Fusarium graminearum-*induced shoot elongation is positively correlated with *F. graminearum* seedling blight severity

To investigate the relationship between *F. graminearum*-induced growth responses and *F. graminearum* seedling blight severity, linear regression analysis was performed on the extent of these changes and seedling survival. This showed that *F. graminearum*-induced seedling elongation has a significant negative correlation with seedling survival rate at 14 days after inoculation (Figure 6A; Pearson’s R^2^ = 0.244; p < 0.005). Thus, stronger fungus*-*induced shoot elongation may be predictive of more severe *F. graminearum* seedling blight. Unexpectedly, the extents of *F. graminearum*-induced shoot elongation showed strong positive correlation with root length (Figure 6B; Pearson’s R^2^ = 0.676; p < 0.005), and a significant negative correlation between root reduction and seedling survival rate (Figure 6C; Pearson’s R^2^ = 0.263; p < 0.005). This leads to the counter-intuitive inference that resistance to fungus-induced root growth inhibition co-occurs with stronger shoot elongation and more severe *F. graminearum* seedling blight.

**Figure 6.**
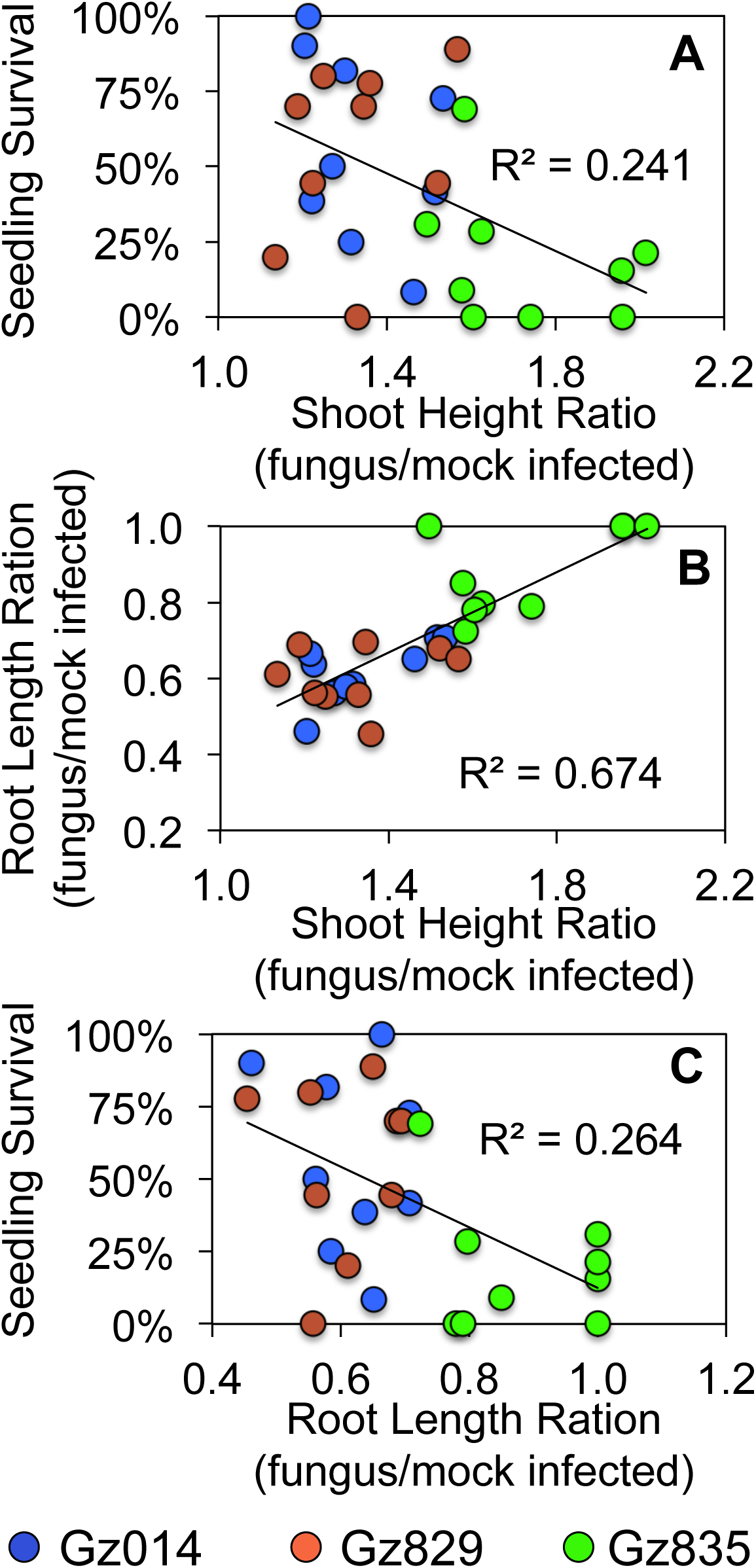
*Fusarium graminearum*-. -induced growth parameter changes in roots and shoots, and seedling blight severity, which are significantly correlated with each other. Each mark represents a maize hybrid-fungal isolate pair, with different fungal isolates represented by different colors. Root length and seedling height were measured at 9 days after infection. Survival was assessed at 14 days after infection

## DISCUSSION

Under field conditions, maize seedlings infected by *F. graminearum* as well as other fungal and oomycete pathogens are often characterized by stunted growth and dehydration in the aboveground tissues before death. Similar seedling stunting symptoms have also been observed in a previous artificial inoculation test with *F. graminearum* spores under laboratory conditions (Kuhnem et al., 2015). However, in our concentration gradient experiment with the same fungal isolates, low dosage *F. graminearum* inoculation on maize seedling root induced shoot elongation (Figure 1A; Figure 2A). Based on the natural variation in this *F. graminearum*-induced response across different maize inbred lines and hybrids reported in this study (Figure 3A; Figure 4A), it is likely that the inconsistent observations between studies could arise from inoculum concentration, maize genotype, or unintended differences between experimental conditions.

Interestingly, the observed *F. graminearum-*induced shoot elongation phenotype is reminiscent of the foolish seedling disease in rice, which is caused by a related fungal pathogen *Gibberella fujikuroi*. Rice seedlings affected by foolish seedling disease demonstrate abnormal seedling elongation resulting from the production of gibberellic acid (GA), a growth-promoting phytohormone, *G. fujikuroi* (Yabuta and Sumiki 1938). Fungus-derived GA has been hypothesized to be a virulence factor by negatively influencing the signaling transduction pathway of the immunity-regulating phytohormone, jasmonic acid (Navarro et al. 2008). However, unlike *G. fujikuroi, F. graminearum* is not known to produce GA, and its genome does not contain biosynthetic genes required for GA biosynthesis (Cuomo et al. 2007). However, we cannot rule out the possibility that *F. graminearum* infection induces endogenous maize GA production.

Consistent with previous reports (Ye et al. 2013; Kuhnem et al., 2015), *F. graminearum* infection induced maize seedling root growth reduction in a host genotype-dependent manner (Figure 1B; Figure 2B; Figure 3B; Figure 4B). Application of a computer image-processing tool, RootReader 2D (Clark et al. 2013), in our study enabled more accurate, quantitative, and efficient measurement of fungus-induced growth response, and its natural variation with host and pathogen genotype. Fungus-induced root reduction and shoot elongation measurements from the same experiment on fifteen maize inbred lines showed no correlation in the extent of these two traits, with some inbred lines showing only one or the other fungus-induced change (Figure 3A&B). This observation suggests that these two responses may be governed by distinct physiological mechanisms.

As one of the most widespread fungal phytopathogens, *F. graminearum* is known to be highly diverse, and different strains are highly variable in their aggressiveness (Backhouse 2014; Boutigny et al. 2011; Lee et al. 2010; Sampietro et al. 2011; Wang et al. 2011). Our survey of nine commercial maize hybrids and three fungal isolates (Figure 4 & 5) is consistent with prior observations that the prevalence of *F. graminearum*-related diseases is highly variable across multiple locations, depending on the aggressiveness of the predominant isolate and environmental permissiveness (Backhouse 2014; Boutigny et al. 2011; Lee et al. 2010; Sampietro et al. 2011; Wang et al. 2011).

Unexpectedly, maize hybrid-fungal isolate pairs with stronger fungus-induced root growth inhibition tended to show a lower level of induced shoot elongation (Figure 6C) and better survival (Figure 6B). This was surprising, because severe reduction in root growth was expected to significantly impair the seedlings’ ability to acquire water and nutrients from the soil, which could lead to seedling death. Two-way ANOVA and a closer look at the data distribution suggest that these correlative relationships are primarily driven by differences between the fungus isolates. Specifically, the isolate that induced significantly stronger shoot elongation, Gz835, induced less root reduction and seedling death (Figure 5). This suggests that these fungus-induced growth responses can be used for quantitative assessment of the aggressiveness of *F. graminearum* isolates, and that different *F. graminearum* isolates may have distinct pathogenic mechanisms when interacting with maize seedlings. Further research with maize seedlings growing in agricultural fields will be required to determine whether *F. graminearum*-induced shoot elongation can serve as a reliable predictor of fungal sensitivity in commercially grown maize.

## ACKNOWLEDGEMENT

The authors would like to thank the Blue River Organic Seed Company for providing the maize hybrids used in this study, Jon Shaff and Eric Craft for training and assistance in using the RootReader 2D platform, and Jaime Cummings for training and assistance in fungal culture maintenance. This work was funded by the Northeast Sustainable Agriculture and Research Education Graduate Student Grant to S.Z. (GNE14-092), USDA-NIFA Hatch Grant NYC1537436 to G.C.B., and US National Science Foundation IOS-1339237 to G.J. J.S.B. was supported by the Plant Genome Research Program at the Boyce Thompson Institute, a Research Experience for Undergraduate program funded by the US National Science Foundation award DBI-1358843.

## Author Contribution

S.Z. conceived, designed, and performed the experiments, analyzed the data, and wrote the manuscript. J.S.B. assisted in the natural variation experiments and related data analyses. G.C.B. provided the fungal isolates used in the experiments, and advice in experimental design and manuscript revision. G.J. provided infrastructural support to this study, and advice in experimental design and manuscript revision.

## References

Ali ML, Taylor JH, Jie L, Sun G, William M, Kasha KJ, Reid LM, Pauls KP (2005) Molecular mapping of QTLs for resistance to Gibberella ear rot, in corn, caused by Fusarium graminearum. Genome 48 (3):521–533. doi:10.1139/g05-014

Arunachalam C, Doohan FM (2013) Trichothecene toxicity in eukaryotes: Cellular and molecular mechanisms in plants and animals. Toxicol Lett 217 (2):149– 158. doi:10.1016/j.toxlet.2012.12.003

Backhouse D (2014) Global distribution of Fusarium graminearum, F. asiaticum and F. boothii from wheat in relation to climate. Eur J Plant Pathol 139 (1):161– 173. doi:10.1007/s10658-013-0374-5

Boutigny AL, Ward TJ, Van Coller GJ, Flett B, Lamprecht SC, O’Donnell K, Viljoen A (2011) Analysis of the Fusarium graminearum species complex from wheat, barley and maize in South Africa provides evidence of species-specific differences in host preference. Fungal Genet Biol 48 (9):914–920. doi:10.1016/j.fgb.2011.05.005

Brauner PC, Melchinger AE, Schrag TA, Utz HF, Schipprack W, Kessel B, Ouzunova M, Miedaner T (2017) Low validation rate of quantitative trait loci for Gibberella ear rot resistance in European maize. Theor Appl Genet 130 (1):175–186. doi:10.1007/s00122-016-2802-3

Chen Q, Song J, Du WP, Xu LY, Jiang Y, Zhang J, Xiang XL, Yu GR (2017) Identification, mapping, and molecular marker development for Rgsr8.1: a new quantitative trait locus conferring resistance to Gibberella stalk rot in maize (Zea mays L.). Front Plant Sci 8:1355. doi:10.3389/fpls.2017.01355

Clark RT, Famoso AN, Zhao KY, Shaff JE, Craft EJ, Bustamante CD, Mccouch SR, Aneshansley DJ, Kochian LV (2013) High-throughput two-dimensional root system phenotyping platform facilitates genetic analysis of root growth and development. Plant Cell Environ 36 (2):454–466. doi:10.1111/j.1365-3040.2012.02587.x

Cuomo CA, Guldener U, Xu JR, Trail F, Turgeon BG, Di Pietro A, Walton JD, Ma LJ, Baker SE, Rep M, Adam G, Antoniw J, Baldwin T, Calvo S, Chang YL, Decaprio D, Gale LR, Gnerre S, Goswami RS, Hammond-Kosack K, Harris LJ, Hilburn K, Kennell JC, Kroken S, Magnuson JK, Mannhaupt G, Mauceli E, Mewes HW, Mitterbauer R, Muehlbauer G, Munsterkotter M, Nelson D, O’Donnell K, Ouellet T, Qi W, Quesneville H, Roncero MI, Seong KY, Tetko IV, Urban M, Waalwijk C, Ward TJ, Yao J, Birren BW, Kistler HC (2007) The Fusarium graminearum genome reveals a link between localized polymorphism and pathogen specialization. Science 317 (5843):1400–1402. doi:10.1126/science.1143708

Kebede AZ, Woldemariam T, Reid LM, Harris LJ (2016) Quantitative trait loci mapping for Gibberella ear rot resistance and associated agronomic traits using genotyping-by-sequencing in maize. Theor Appl Genet 129 (1):17–29. doi:10.1007/s00122-015-2600-3

Kuhnem PR, Del Ponte EM, Dong YH, Bergstrom GC (2015) Fusarium graminearum ssolates from wheat and maize in New York show similar range of aggressiveness and toxigenicity in cross-species pathogenicity tests. Phytopathology 105 (4):441–448. doi:10.1094/Phyto-07-14-0208-R

Lee SH, Lee J, Nam YJ, Lee S, Ryu JG, Lee T (2010) Population structure of Fusarium graminearum from maize and rice in 2009 in Korea. Plant Pathology J 26 (4):321–327. doi:10.5423/Ppj.2010.26.4.321

Ma C, Ma X, Yao L, Liu Y, Du F, Yang X, Xu M (2017) qRfg3, a novel quantitative resistance locus against Gibberella stalk rot in maize. Theor Appl Genet 130 (8):1723–1734. doi:10.1007/s00122-017-2921-5

Maresca M (2013) From the gut to the brain: journey and pathophysiological effects of the food-associated trichothecene mycotoxin deoxynivalenol. Toxins 5 (4):784–820. doi:10.3390/toxins5040784

McMullen M, Bergstrom G, De Wolf E, Dill-Macky R, Hershman D, Shaner G, Van Sanford D (2012) A unified effort to fight an enemy of wheat and barley: Fusarium head blight. Plant Dis 96 (12):1712–1728. doi:10.1094/Pdis-03-12-0291-Fe

McMullen MD, Kresovich S, Villeda HS, Bradbury P, Li H, Sun Q, Flint-Garcia S, Thornsberry J, Acharya C, Bottoms C, Brown P, Browne C, Eller M, Guill K, Harjes C, Kroon D, Lepak N, Mitchell SE, Peterson B, Pressoir G, Romero S, Oropeza Rosas M, Salvo S, Yates H, Hanson M, Jones E, Smith S, Glaubitz JC, Goodman M, Ware D, Holland JB, Buckler ES (2009) Genetic properties of the maize nested association mapping population. Science 325 (5941):737–740. doi:10.1126/science.1174320

Mueller D (2016a) Corn disease loss estimates from the United States and Ontario, Canada - 2013. Purdue University Extension, Retrieved from http://cropprotectionnetwork.org/crop-loss-estimates/corn-disease-loss-estimates-2013/.

Mueller D (2016b) Corn disease loss estimates from the United States and Ontario, Canada - 2014. Purdue University Extension, Retrieved from http://cropprotectionnetwork.org/crop-loss-estimates/corn-disease-loss-estimates-2014/.

Mueller D (2016c) Corn disease loss estimates from the United States and Ontario, Canada - 2015. Purdue University Extension, Retrieved from http://cropprotectionnetwork.org/crop-loss-estimates/corn-disease-loss-estimates-2015/.

Mueller D (2017) Corn disease loss estimates from the United States and Ontario, Canada - 2016. Purdue University Extension, Retrieved from http://cropprotectionnetwork.org/crop-loss-estimates/corn-disease-loss-estimates-2016/.

Munkvold GP, White DG (2016) Compendium of Corn Diseases. Fourth edn. The American Phytopathological Society,

Navarro L, Bari R, Achard P, Lison P, Nemri A, Harberd NP, Jones JDG (2008) DELLAs control plant immune responses by modulating the balance and salicylic acid signaling. Curr Biol 18 (9):650–655. doi:10.1016/j.cub.2008.03.060

Sampietro DA, Diaz CG, Gonzalez V, Vattuone MA, Ploper LD, Catalan CAN, Ward TJ (2011) Species diversity and toxigenic potential of Fusarium graminearum complex isolates from maize fields in northwest Argentina. Int J Food Microbiol 145 (1):359–364. doi:10.1016/j.ijfoodmicro.2010.12.021

Trail F (2009) For Blighted Waves of Grain: Fusarium graminearum in the Postgenomics Era. Plant Physiol 149 (1):103–110. doi:10.1104/pp.108.129684

Wang JH, Ndoye M, Zhang JB, Li HP, Liao YC (2011) Population structure and Genetic diversity of the Fusarium graminearum species complex. Toxins 3 (8):1020– 1037. doi:10.3390/toxins3081020

Yabuta T, Sumiki Y (1938) On the crystal of gibberellin, a substance to promote plant growth. Journal of the Agricultural Chemical Society of Japan 14 (1526)

Yang Q, Yin GM, Guo YL, Zhang DF, Chen SJ, Xu ML (2010) A major QTL for resistance to Gibberella stalk rot in maize. Theor Appl Genet 121 (4):673–687. doi:10.1007/s00122-010-1339-0

Ye JR, Guo YL, Zhang DF, Zhang N, Wang C, Xu ML (2013) Cytological and molecular characterization of quantitative trait locus qRfg1, which confers resistance to Gibberella stalk rot in maize. Mol Plant Microbe In 26 (12):1417–1428. doi:10.1094/Mpmi-06-13-0161-R

Zhang DF, Liu YJ, Guo YL, Yang Q, Ye JR, Chen SJ, Xu ML (2012) Fine-mapping of qRfg2, a QTL for resistance to Gibberella stalk rot in maize. Theor Appl Genet 124 (3):585–596. doi:10.1007/s00122-011-1731-4

Zhang Y, He J, Jia LJ, Yuan TL, Zhang D, Guo Y, Wang Y, Tang WH (2016) Cellular Tracking and gene profiling of Fusarium graminearum during maize stalk rot disease development elucidates its strategies in confronting phosphorus limitation in the host apoplast. PLoS Pathog 12 (3):e1005485. doi:10.1371/journal.ppat.1005485

